# An ecological niche model to predict the geographic distribution of *Haemagogus janthinomys*, Dyar, 1921 the yellow fever and Mayaro virus vector, in South America

**DOI:** 10.1101/2021.04.05.437767

**Authors:** Michael Celone, David Pecor, Alexander Potter, Alec Richardson, James Dunford, Simon Pollet

**Affiliations:** Uniformed Services University of Health Sciences, Bethesda, Maryland, United States of America; Department of Entomology, Walter Reed Army Institute of Research, Silver Spring, Maryland, United States of America; Walter Reed Biosystematics Unit, Suitland, Maryland, United States of America; Infectious Disease Clinical Research Program, Department of Preventive Medicine and Biostatistics, Uniformed Services University of the Health Sciences, Bethesda, Maryland, United States of America; Henry M. Jackson Foundation for the Advancement of Military Medicine, Inc., Bethesda, Maryland, United States of America

## Abstract

Yellow fever virus (YFV) has a long history of impacting human health in South America. Mayaro virus (MAYV) is an emerging arbovirus of public health concern in the Neotropics and its full impact is yet unknown. Both YFV and MAYV are primarily maintained via a sylvatic transmission cycle but can be opportunistically transmitted to humans by the bites of infected forest dwelling *Haemagogus janthinomys* Dyar, 1921. To better understand the potential risk of YFV and MAYV transmission to humans, a more detailed understanding of this vector species’ distribution is critical. This study compiled a comprehensive database of 170 unique *Hg. janthinomys* collection sites retrieved from the published literature, digitized museum specimens and publicly accessible mosquito surveillance data. Covariate analysis was performed to optimize a selection of environmental (topographic and bioclimatic) variables associated with predicting habitat suitability, and species distributions modelled across South America using a maximum entropy (MaxEnt) approach. Our results indicate that suitable habitat for *Hg. janthinomys* can be found across forested regions of South America including the Atlantic forests and interior Amazon.

**Author Summary:** Mayaro virus is a neglected tropical disease and there is insufficient evidence to define its geographic range. The mosquito *Haemagogus janthinomys* is a primary vector of Mayaro and its distribution is largely unknown at a sub-country scale. Building compendiums of collection data and creating ecological niche models provides a more precise estimation vector species potential habitat. Our dataset stands as one of the most expansive existing for collection data of this species combining data published in literature, publicly available data repositories and digitized museum specimen records. Comparing results of niche models with near real time environmental data can give even better predictions of areas where Mayaro virus exposure could occur. The methods and results of this study can be replicated for any disease/vector of interest so long as there is data discoverable through the scientific literature, public repositories, or other civilian and governmental agencies willing to share.

## Introduction

Yellow Fever virus (YFV) is a mosquito-borne flavivirus that causes symptoms including fever, muscle pain, nausea, and fatigue. Although many people recover from initial symptoms of yellow fever, approximately 15 percent of infected patients experience more severe infections including hemorrhage, jaundice, and damage to multiple organ systems (1), with case fatality rates exceeding 40% (2). Globally, approximately 400 million people are estimated to be at risk of YFV (3). Although widespread vaccination campaigns have reduced the burden of YFV circulation, several epidemics and epizootics have occurred in South America during the last two decades, primarily in Brazil (4). In the Americas, YFV predominantly circulates in a sylvatic transmission cycle involving non-human primates and canopy-dwelling mosquitoes of the *Haemagogus* subgenus *Haemagogus* (5). Contemporary human YFV outbreaks represent spillover events from this sylvatic cycle into the human population (6).

Mayaro virus (MAYV) is a recently emerging arbovirus with a sylvatic transmission cycle throughout Central and South America that occasionally spills over into human populations in Brazil, Bolivia, and Venezuela (7). While MAYV is not known to be fatal, it can cause non-specific febrile symptoms, and occasionally results in debilitating polyarthritis or polyarthralgia (8). Although the precise burden of MAYV is unknown, human seroprevalence surveys have detected MAYV circulation in many countries including Peru (9), Suriname (10), Mexico (11), Colombia (12), French Guiana (13), and Haiti (14). Canopy-dwelling, *Haemagogus janthinomys* Dyar, 1921 is among several mosquito species that are considered important vectors of both YFV and MAYV (15).

A comprehensive understanding of the geographic distribution of *Hg. janthinomys* mosquitoes is essential to predicting areas at risk of MAYV and YFV outbreaks. However, it is impossible to exhaustively survey this species across its entire range. Knowledge of the ecological niche preferences of this species can guide disease surveillance efforts and aid public health authorities in allocating resources for vector control measures. Ecological niche modeling (ENM) techniques have been used extensively to predict the potential range of disease vectors (16), including vectors of Rift Valley fever virus (17), *Trypanosoma cruzi* (18) and Japanese encephalitis virus (19). Although recent modeling studies have used ENM frameworks such as the maximum entropy (MaxEnt) approach to predict the geographic range of the Mayaro (20) and yellow fever viruses (21), there have been no prior attempts to model the habitat suitability of the major vector, *Hg. janthinomys*. This study aims to develop a robust species distribution model of *Hg. janthinomys* using a comprehensive dataset of collection records compiled from publicly accessible databases and peer-reviewed literature.

## Methods

Distribution data for *Hg. janthinomys* were compiled from publicly available specimen collection records, archive specimens in the United States National Museum (USNM) mosquito collection, and records reported in peer-reviewed scientific literature. A search of the VectorMap data repository (vectormap.si.edu) yielded both collection locations from USNM specimen records and mosquito surveillance data. Additional collection events were identified from the Global Biodiversity Information Facility (GBIF) database (22) and the NCBI GenBank database (23).

A literature search was conducted using PubMed, Web of Science and Google Scholar. Searches were executed using the keywords “*Haemagogus janthinomys*” combined with each country in South America, including Trinidad and Tobago, for all articles published between 1901 and December 20, 2020. Trinidad and Tobago were included as it listed as the type locality of *Hg. janthinomys*. The search scope was modified to exclude Central American countries after initial searches yielded very few collection records from the literature.

Articles were considered for eligibility based on the following criteria: (i) original research studies on arthropod vectors in South America that described field-collected *Hg. janthinomys* adult mosquitoes, larvae, or pupae or original research studies that described the bionomics of *Hg. janthinomys*; and (ii) studies that included mappable collection sites (either GPS coordinates or specific named places that can be georeferenced). Articles were not included if they met any of the following exclusion criteria: (i) studies involving only humans; (ii) studies not reporting original data (*e*.*g*., review articles, perspective pieces, editorials, recommendations, and guidelines); (iii) duplicate studies; (iv) laboratory-based vector competence studies or studies involving laboratory-reared mosquitoes; (v) studies that did not provide exact collection site locations; (vi) studies that did not provide information on mosquito identification methods.

All articles were organized using EndNote, and data was abstracted into a Microsoft Excel table. A primary reviewer independently screened all titles and abstracts to determine articles that could immediately be discarded and articles to be included in the second stage of review. During this second stage of review, full text articles were reviewed to identify candidates for inclusion in the study. A secondary reviewer examined the screening results to verify the final list of eligible articles. From those studies that met our inclusion criteria, collection data were extracted focusing on all information relevant to preserving the collection event (24). Locality data for each collection event was georeferenced using the point-radius method (25) and data was standardized using the WRBU/VectorMap Best Practices Guide to Data Management and Reporting (26). **See S1 Table** for a full list of sources and coordinates for each collection location.

The MaxEnt approach to developing ENMs has emerged as one of the most popular techniques for species distribution modeling due to its high predictive accuracy (27). This technique uses presence-only collection data and a set of 26 environmental (topographic and bioclimatic) covariates, contrasting environmental conditions at presence points against randomly selected background (test) points (28). Comparative modeling studies have demonstrated that MaxEnt performs better than similar species distribution modeling procedures (*e*.*g*., BioClim, GARP, and Domain) (29, 30).

A total of 26 topographic, climate, and landscape variables were considered for inclusion in the model. The 19 bioclimatic variables from the WorldClim version 2 website were downloaded at a 30-second (1-km) spatial resolution (31). The Global Multi-resolution Terrain Elevation Data (GMTED) slope and elevation datasets were downloaded from the ESRI Living Atlas of the World database at a 7.5 arc-second (∼250 meter) spatial resolution (32). A land use/land cover dataset from MDAUS BaseVue 2013 was also downloaded from the ESRI Living Atlas of the World at a 30-m spatial resolution (33). The Food and Agricultural Organization’s Digital Soil Map of the World was downloaded at a spatial resolution of 5 arc-minutes (34). Additional raster layers were created from a digital elevation map (DEM) raster using the flow direction, flow accumulation, and aspect tools from the ESRI ArcGIS Pro Spatial Analyst toolbox (35). All raster layers were clipped to the extent of South America (15.925 °N, 55.983 °S, 109.458 °W, 28.841 °E) and re-sampled to a 1-km resolution gridded longitude-latitude coordinate system using ArcGIS Pro, resulting in 9674 longitude x 8629 latitude pixels, with a grid-cell resolution of 0.00833 deg. Country shapefiles were accessed through the geoBoundaries Global Administrative Database (36).

## Analysis

A covariate significance assessment was conducted on the sample record collection dataset in order to develop a more refined, statistically robust ENM (37). Covariate significance was assessed via the t-test and r^2^-maximization criteria at the 95% confidence level (α=0.05). Number density was used as the dependent variable, and three multiple least-squares regression models were developed and tested to evaluate the linear and quadratic terms of each covariate: 1) a linear-only (LO) model using the linear terms of each covariate; 2) a quadratic-only (QO) model using the quadratic terms of each covariate; and 3) a linear-quadratic (LQ) model that incorporates both the linear and quadratic terms of each covariate.

The Multiple Addition (MA) and Multiple Removal (MR) methods were used to sequentially add and remove, respectively, one covariate at a time to and from the developing regression model, and to assess the statistical significance of the covariate addition/removal at each successive step. In the MA process, covariates were sequentially added in the order of highest to lowest significance until all significant (P < 0.05) covariates were added. In the MR process, covariates were sequentially removed in the order of lowest to highest significance until all insignificant (P > 0.05) covariates were removed (37). A total of 6 optimal model runs were conducted: three models (LO, QO, LQ) by two sequential methods (MA, MR). A covariate was labeled as significant if at least one of its terms (linear or quadratic) was retained in at least one of the six optimal models (to establish consensus among models).

A Comparison of Regressions analysis was conducted between the Full LO and QO regression models (retaining all linear or quadratic covariate terms, respectively) to assess differences in correlation coefficient (r), regression slopes (between the linear and quadratic terms of each covariate), elevation, and coincidental regression between these two models. A significant difference between the linear and quadratic slopes for a given covariate would indicate significant non-linear dependence of the dependent response variable (*i*.*e*., number density) on that covariate. Bilinear interpolation was used to estimate covariate values at each sample record collection data point by overlaying the covariate raster grids on the sample data points, identifying the 1-km resolution grid-box that each sample data point resides in, converting the geographic coordinates of the sample data point and four corner points of the grid-box to easting-northing, and calculating spatial distances as the basis for interpolation.

A fundamental assumption of ENMs is that species occurrence records are collected through systematic or random sampling (*i*.*e*., unbiased samples) (28), but this assumption is often violated when certain areas are oversampled because they are more easily accessible (38). This spatial bias can reduce model accuracy because environmental features of these more accessible areas are overrepresented in the model, leading to issues like spatial autocorrelation and errors of omission or commission (39). One solution to correct for sampling bias is the selection of background points with the same selection bias as the presence points (38). MaxEnt’s default procedure is to select background points at random from the study extent (28). However, we generated a ‘bias file’ using the *MASS* package in the R statistical software to ensure that the background sampling represented the record density of the occurrence points. As a result, background points were chosen preferentially from areas with a high density of presences.

MaxEnt version 3.4.1 was used to generate a distribution map of *Hg. janthinomys* habitat suitability using topographic, landscape, and climate variables. An output format of *cloglog* was selected, which returns an estimate between 0 and 1 that represents the probability of presence within each pixel (40). The model was run using the *k-fold* cross validation replicated run type with 10 replicates. For this validation procedure, the data set is split into *k* independent subsets (*i*.*e*., folds), and the model is trained on *k-1* subsets and validated using the k^th^ subset (28). Therefore, each replicate in our model was trained on approximately 90% of the occurrence records and validated using approximately 10% of the occurrence data. In addition, the maximum iterations were set to 10,000 and all other settings were left as their defaults. Features were selected automatically based on the findings of a similar study that reported improved model performance with automatic feature selection versus manual selection (41). The jackknife test as well as the percent contribution and permutation importance were used to assess the relative importance of each variable in the model. In order to perform the jackknife procedure, the MaxEnt program runs several models where each variable is omitted in turn, followed by additional models where each variable is used in isolation to predict the species distribution.

An initial model was run with all 26 variables and a reduced model was run with the five significant variables identified from the covariate analysis. The area under the receiver-operating curve (AUC) was used to assess model performance, based on the average AUC across the 10 model replicates. An AUC of 1 suggests that the model perfectly predicts the distribution of the vector while an AUC of 0.5 suggests that the model is not able to predict the distribution better than random chance. We used the AUC of test data (AUC_TEST_) to assess model performance instead of the AUC of training data (AUC_TRAIN_) due to the problems of overfitting associated with the AUC_TRAIN_ statistic (42).

## Results

Overall, 170 unique geolocations of verified *Hg. janthinomys* presence from 11 countries in South America (including Trinidad and Tobago) were used to build the ENM, with maximal records from Brazil (n=110) and French Guiana (n=23) (**see Fig 1 and Table 1**). Five covariates were selected for inclusion in the final ENM based on the covariate significance analysis: one categorical (land cover) and four continuous (slope, Mean Temperature of Wettest Quarter, Precipitation Seasonality, Precipitation of Warmest Quarter) (**See Table 2 for variable descriptions**).

**Table 1.** Summary of *Hg. janthinomys* presence points by country.

| Country | Total (n) | Total (%) |
| --- | --- | --- |
| Brazil | 110 | 64.7% |
| French Guiana | 23 | 13.5% |
| Colombia | 11 | 6.5% |
| Trinidad & Tobago | 9 | 5.3% |
| Venezuela | 8 | 4.7% |
| Ecuador | 3 | 1.8% |
| Suriname | 2 | 1.2% |
| Argentina | 1 | 0.6% |
| Bolivia | 1 | 0.6% |
| Guyana | 1 | 0.6% |
| Peru | 1 | 0.6% |
| **Total** | **170** | **100.0%** |

**Table 2.** **Minimum, maximum, average values, percent contribution, and permutation importance of variables in the *Hg. janthinomys* model**

| Variable | Description | Min. | Max. | Avg. | Contribution (%) | Permutation (%) |
| --- | --- | --- | --- | --- | --- | --- |
| **Slope** | Values represent the angle of the downward sloping terrain from 0 (flat) to 90 degrees (vertical) | 0 | 33 | 4.9 | 9 | 31.4 |
| **LU/LC** | Categorical variable with 16 classes of land use / land cover | N/A<sup>a</sup> | N/A | N/A | 76.9 | 27 |
| **BIO8** | Mean Temperature of Wettest Quarter | 19.6 | 27.5 | 24.8 | 4 | 33.7 |
| **BIO15** | Precipitation Seasonality (Coefficient of Variation), mm | 10.2 | 114.8 | 55.5 | 7.6 | 4.5 |
| **BIO18** | Precipitation of Warmest Quarter | 67 | 869 | 421.1 | 2.6 | 3.4 |

<sup>a</sup> Minimum, maximum, and average values were not calculated for the categorical variable land use/land cover.

**Figure 1.**
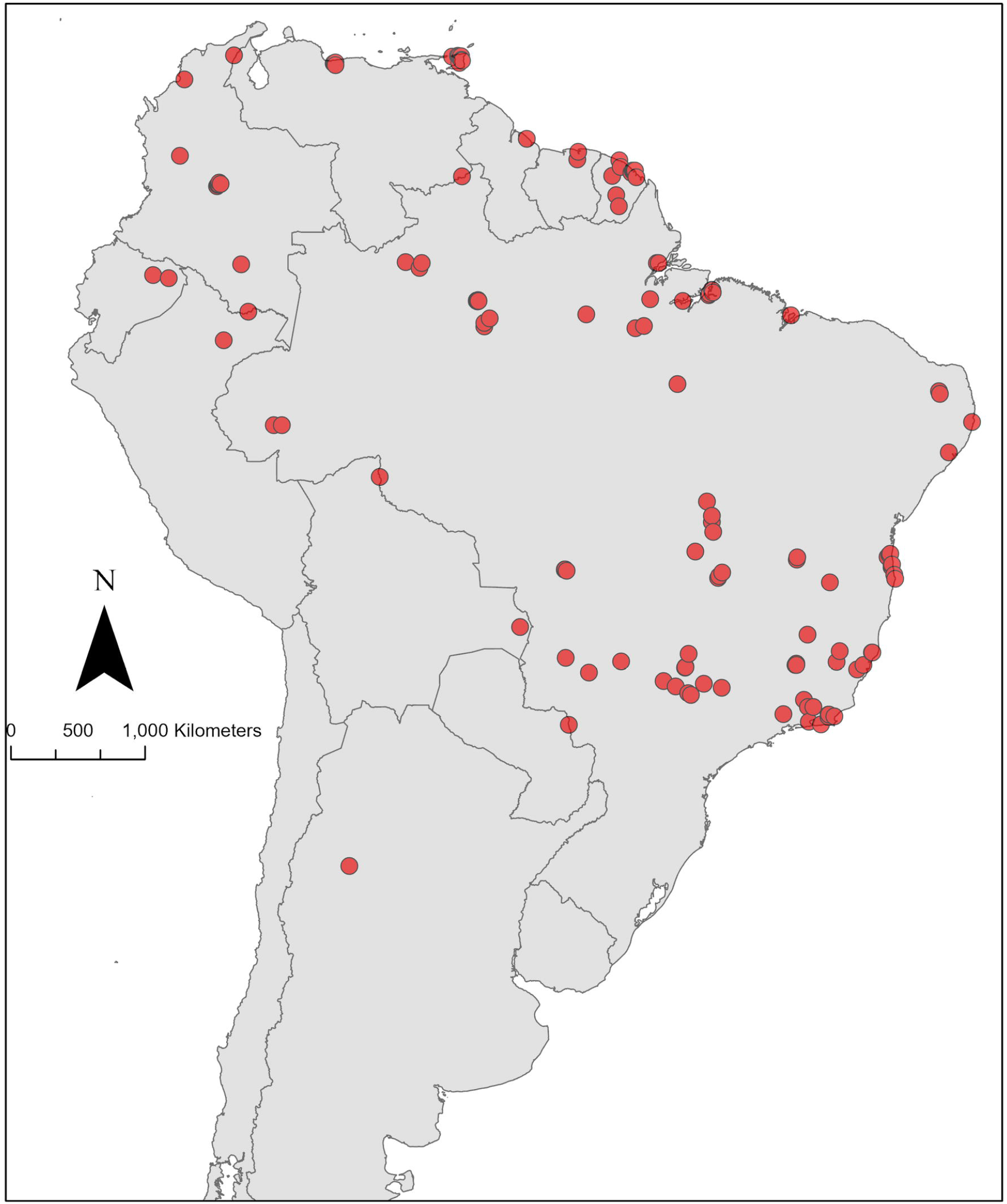
Map of *Hg. janthinomys* presence locations. Points represent locations where *Hg*. janthinomys mosquitoes were collected.

The habitat suitability map for *Hg. janthinomys* based on both the reduced and full models are presented in **Fig 2**. The reduced suitability map (**Fig 2A**) represents the average of the 10 replicate runs incorporating the five most significant variables identified by the covariate significance assessment. The average area under the receiver operating characteristic curve for testing data (AUC_TEST_) across the 10 model replicates was 0.75 (SD±0.07). The analysis of variable contribution demonstrated that land use/land cover contributed the greatest amount of information to the model (76.9%), followed by slope (9%). Similarly, the jackknife tests of both training gain and test gain revealed that land use/land cover was the most important variable for developing the ENM. In other words, this variable increased the training/test gain most substantially when used in isolation and decreased the training/test gain most substantially when omitted from the model. In addition, the permutation importance was greatest for BIO8-*mean temperature of wettest quarter* (33.7%) followed by slope (31.4%) and land use/land cover (27.0%). The minimum, maximum, and average values of the four continuous variables were extracted at each *Hg. janthinomys* collection point. The results are presented in **Table 2** along with the percent contribution and permutation importance. The response curves are presented in **Fig 3**.

**Figure 2.**
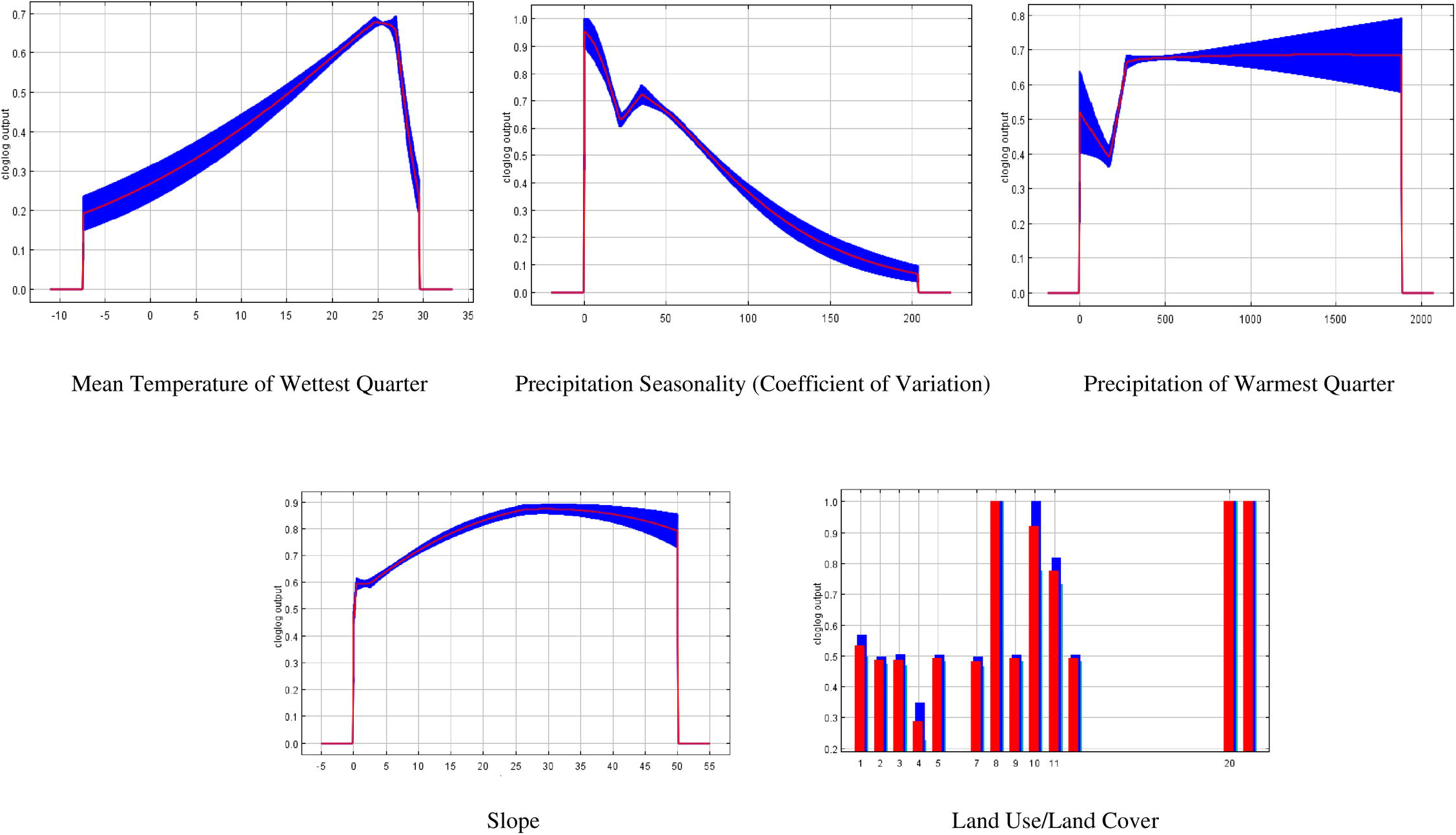
Estimated distribution of the probability of *Hg. janthinomys* presence according to the MaxEnt models: A) reduced model, and B) full model. Red represents areas of highest suitability for *Hg. janthinomys* while blue represents areas of low suitability.

**Figure 3.**
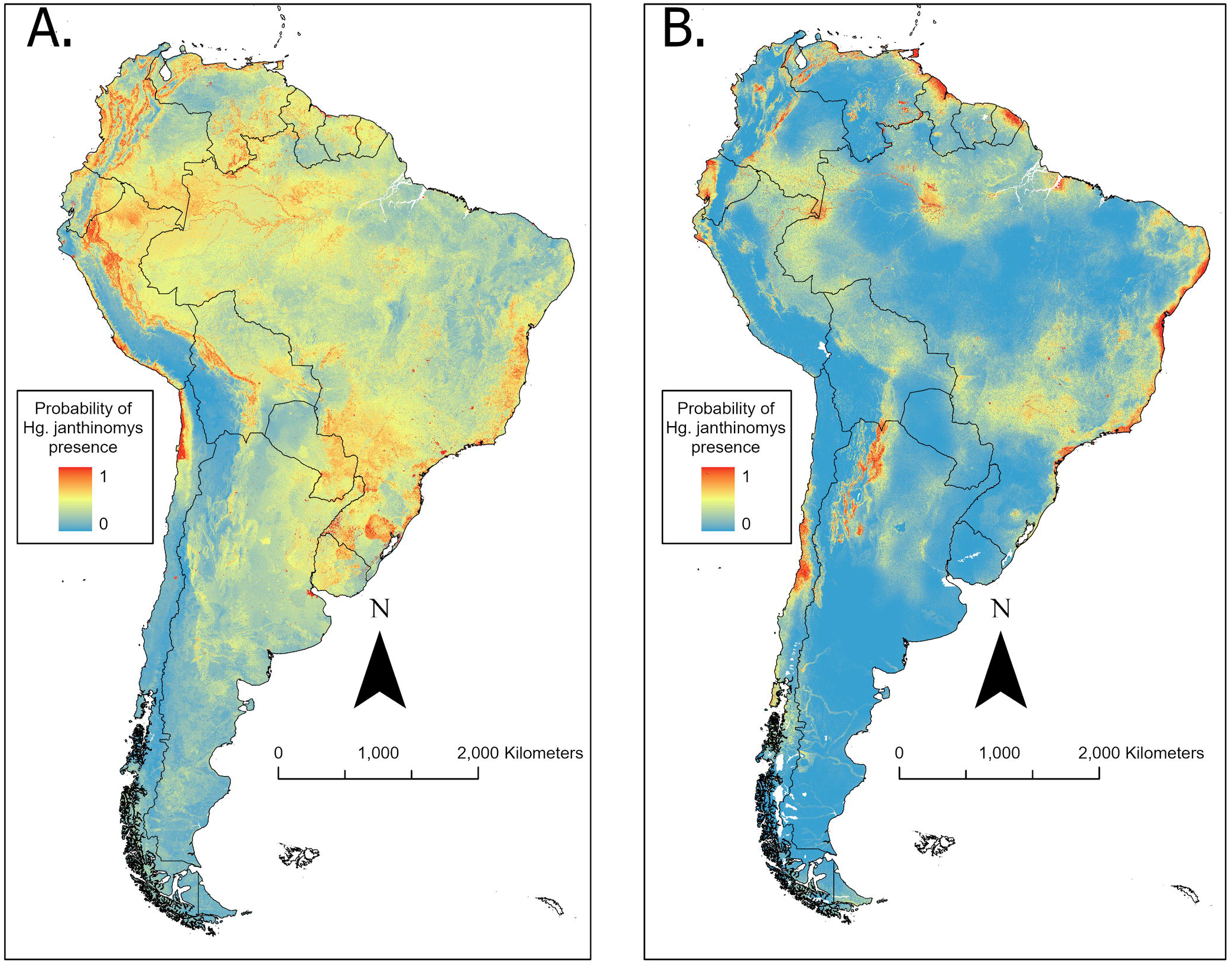
Response curves for the reduced *Hg. janthinomys* model. Each curve represents a Maxent model created using only the corresponding variable. The red lines represent the mean response of 10 Maxent runs while blue represents the mean ± 1 standard deviation. Land cover classes are: (1) Deciduous forest; (2) Evergreen forest; (3) Shrub/Scrub; (4) Grassland; (5) Barren or minimal vegetation; (7) Agricultural, General; (8) Agriculture, Paddy; (9) Wetland; (10) Mangrove; (11) Water; (12) Ice/Snow; (20) High Density Urban; (21) Medium-Low Density Urban.

The full suitability map (**Fig 2B**) represents the average of the 10 replicate runs incorporating all 26 bioclimatic variables. The average area under the receiver operating characteristic curve for testing data (AUC_TEST_) across the 10 model replicates was 0.87 (SD±0.04). The analysis of variable contribution demonstrated that land use/land cover contributed the greatest amount of information to the model (32.4%), followed by BIO2-*mean diurnal range* (25.3%) and BIO5-*maximum temperature of warmest month* (7.6%). Similarly, the jackknife tests of both training gain and test gain revealed that land use/land cover and BIO2 were the most important variables for developing the ENM. In addition, the permutation importance was greatest for BIO4-*temperature seasonality* (13.4%) followed by BIO2-*mean diurnal range* (10.5%). The minimum, maximum, and average values of the continuous variables were extracted at each *Hg. janthinomys* collection point. The results are presented in the **S2 table** along with the percent contribution and permutation importance.

## Discussion

This study has produced a comprehensive database of *Hg. janthinomys* collection events. As an important vector of YF and MAY viruses, it is crucial for health planners to be informed of where this species may contribute to human infections across South America. Our study represents a contemporary prediction of the mosquito’s potential ecological niche, incorporating the publicly available records of *Hg. janthinomys* presence. The model may guide surveillance activities for *Hg. janthinomys* by identifying high vector-presence suitability regions.

ENMs provide the opportunity to predict the locale of a species across a large geographic range using a relatively small number of collection records. Our reduced and full models predicted the potential ecological niche of this mosquito with an AUC of 0.75 and 0.86 respectively, demonstrating several regions of high suitability throughout South America. These findings are consistent with an ecological niche model published in 2010, which found optimal conditions for *Hg. janthinomys* along the coast of northeast Brazil and northern Venezuela, based on 78 presence records (43).

Although MAYV outbreaks have only occurred sporadically, the Pan American Health Organization (PAHO) has emphasized its growing importance and recommended increased surveillance and enhanced diagnostic capacity (44). Our models identified areas of moderate-to-high suitability for *Hg. janthinomys*, the primary MAYV vector, in the central and northern regions of French Guiana, which aligns with recent epidemiological findings. For example, a recent seroprevalence survey found circulation of MAYV throughout French Guiana (45), and the World Health Organization (WHO) recently reported 13 confirmed MAYV cases there over a 3-month period (46). Additional locations of MAYV outbreaks in Para, Brazil (47, 48) and Portuguesa, Venezuela (49) have also occurred in regions of moderate-to-high suitability for the vector according to our models. *Hg. janthinomys* has also been identified as the primary vector during recent outbreaks of sylvatic YFV in Brazil (50). During these outbreaks, the majority of YFV cases occurred in areas of moderate-to-high suitability, especially along the coastal areas in the states of Sao Paulo, Rio de Janeiro, Espirito Santo, and in the eastern portion of Mato Grosso. Therefore, areas predicted as highly suitable for *Hg. janthinomys* occurrence may serve as locations of potential disease spillover that could be targeted for increased surveillance and enhanced vector control and disease mitigation efforts.

Analysis of variable contributions demonstrated that land use/land cover had the greatest relative contribution to our models and the greatest training gain based on the jackknife test. *Hg. janthinomys* is arboreal and has recently been detected in many different forest types including mangrove, semi-evergreen seasonal, evergreen seasonal, and young secondary forest (51). The same study also found *Hg. janthinomys* close to agricultural communities in Trinidad & Tobago and hypothesized that forest fragmentation may play an important role in *Hg. janthinomys* presence (52). Therefore, changing patterns of land use/land cover and encroachment into forested areas may increase human exposure to *Hg. janthinomys* and to the pathogens they transmit. Several studies have suggested an occupational risk to MAYV infection among rainforest hunters (53) and forest crop-plot workers (54). In addition, high MAYV seroprevalence has been found in populations residing close to forested areas (55, 56). Communities in close proximity to the forest should therefore be prioritized for vector and pathogen surveillance to determine if MAYV or YFV are circulating in local mosquito populations.

The response curves for our models also reveal important information about the preferred environmental conditions of the *Hg. janthinomys* mosquito. The most important variable in the reduced model according to the permutation importance was BIO8-*mean temperature of the wettest quarter*. The response curve for this variable demonstrates that the ideal value of BIO8 for mosquito suitability peaks around 26 °C and decreases sharply thereafter. In addition, one of the most important variables in the full model according to permutation importance was BIO2-*mean diurnal range*. The response curve for BIO2 demonstrates that the ideal diurnal range for mosquito suitability peaks at 5 °C and then decreases steadily. Several studies have explored the impact of temperature fluctuations on *Aedes* and *Anopheles* mosquito dynamics (57-59), demonstrating that large diurnal temperature range can affect life history traits including larval development time, adult survival, and reproductive output. Peak activity of *Hg janthinomys* has been found to occur during periods of high temperature (60) and our findings demonstrate that daily temperature fluctuations may also impact habitat suitability for *Hg. janthinomys*.

Our study has several limitations (including sampling bias and a limited set of environmental covariates) that should be considered when interpreting the findings. One major limitation is the sampling bias associated with mosquito collections. Areas of high accessibility, including those in close proximity to roads, are more likely to be sampled, potentially leading to inaccurate models (39). As a result, the presence records that we have compiled do not represent the complete ecological niche of *Hg. janthinomys* but may be biased toward accessible locations. We attempted to correct for sampling bias by including a bias layer during the model-building process, according to techniques proposed by Phillips et al. (38), to ensure selection of background points with the same bias as the presence records. Another limitation inherent in the ecological niche modeling process is the use of a limited set of environmental covariates. While the covariate significance assessment identified insignificant covariates (subsequently removed from the statistically robust ENM) from WorldClim’s complete pool of 26 topographic and bioclimatic covariates, there may be additional covariates not included in this pool that are significant and could be included. Although these variables play an important role in predicting areas of high suitability for *Hg. janthinomys*, other factors such as human population density, socio-economic status, host migration patterns, presence of other mosquito species (in competitive and symbiotic associations for available ecological niches), and intensity of mosquito control and disease mitigation programs can also impact the occurrence probability and geographical distributions of *Hg. janthinomys*. Recommendations for future research may include investigating additional mosquito species, identifying under sampled regions via surveillance gap analysis, conducting follow-up surveillance studies in these under sampled regions to reduce sampling bias, incorporating morphological and molecular data on the MAYV and YFV disease pathogens, and running updated models with the enhanced record collection dataset in efforts to further improve the efficacy and statistical robustness of the ecological niche models.

## Acknowledgments

This research was supported in part by an appointment to the Research Participation Program at the Walter Reed Army Institute of Research by the Oak Ridge Institute for Science and Education through an interagency agreement between the U.S. Department of Energy and USAMRMC. This work was financially supported by the Armed Forces Health Surveillance Division – Global Emerging Infections Surveillance (AFHSB-GEIS) [P0140_20_WR_05] and the Henry M. Jackson Foundation for the Advancement of Military Medicine. The activities undertaken at WRBU were performed in part under a Memorandum of Understanding between the Walter Reed Army Institute of Research (WRAIR) and the Smithsonian Institution, with institutional support provided by both organizations. The Infectious Disease Clinical Research Program is supported by the National Institute of Allergy and Infectious Diseases, National Institute of Health (Inter-Agency Agreement Y1-AI-5072).

## Disclaimer

The view(s) expressed in this article are those of the authors and do not necessarily reflect the official policy or position of the Uniformed Services University of the Health Sciences (USU), Henry M. Jackson Foundation for the Advancement of Military Medicine, Inc., the National Institutes of Health or the Department of Health and Human Services, Departments of the Army, Air Force, or Navy, the Department of Defense, or the U.S. Government. The use of trade names in this document does not constitute an official endorsement or approval of the use of such commercial hardware or software. Do not cite this document for advertisement. The publication has been cleared for publication by the Walter Reed Army Institute of Research (WRAIR) and USU.

## Supporting Information Captions

**S1 Table** Coordinates and citations for locations included in the model.

**S2 Table** Minimum, maximum, average values, percent contribution, and permutation importance of variables in the full *Hg. janthinomys* model.

## References

1. Barnett ED. Yellow fever: epidemiology and prevention. Clinical infectious diseases : an official publication of the Infectious Diseases Society of America. 2007;44(6):850–6.

2. Tuboi SH, Costa ZG, da Costa Vasconcelos PF, Hatch D. Clinical and epidemiological characteristics of yellow fever in Brazil: analysis of reported cases 1998-2002. Transactions of the Royal Society of Tropical Medicine and Hygiene. 2007;101(2):169–75.

3. Shearer FM, Moyes CL, Pigott DM, Brady OJ, Marinho F, Deshpande A, et al. Global yellow fever vaccination coverage from 1970 to 2016: an adjusted retrospective analysis. Lancet Infect Dis. 2017;17(11):1209–17.

4. Monath TP, Vasconcelos PF. Yellow fever. Journal of clinical virology : the official publication of the Pan American Society for Clinical Virology. 2015;64:160–73.

5. Abreu FVS, Ribeiro IP, Ferreira-de-Brito A, Santos A, Miranda RM, Bonelly IS, et al. Haemagogus leucocelaenus and Haemagogus janthinomys are the primary vectors in the major yellow fever outbreak in Brazil, 2016-2018. Emerging microbes & infections. 2019;8(1):218–31.

6. de Oliveira Figueiredo P, Stoffella-Dutra AG, Barbosa Costa G, Silva de Oliveira J, Dourado Amaral C, Duarte Santos J, et al. Re-Emergence of Yellow Fever in Brazil during 2016-2019: Challenges, Lessons Learned, and Perspectives. Viruses. 2020;12(11).

7. Pezzi L, Rodriguez-Morales AJ, Reusken CB, Ribeiro GS, LaBeaud AD, Lourenco-de-Oliveira R, et al. GloPID-R report on chikungunya, o’nyong-nyong and Mayaro virus, part 3: Epidemiological distribution of Mayaro virus. Antiviral research. 2019;172:104610.

8. Suhrbier A, Jaffar-Bandjee MC, Gasque P. Arthritogenic alphaviruses--an overview. Nature reviews Rheumatology. 2012;8(7):420–9.

9. Forshey BM, Guevara C, Laguna-Torres VA, Cespedes M, Vargas J, Gianella A, et al. Arboviral etiologies of acute febrile illnesses in Western South America, 2000-2007. PLoS neglected tropical diseases. 2010;4(8):e787.

10. Jonkers AH, Spence L, Karbaat J. Arbovirus infections in Dutch military personnel stationed in Surinam. Further studies. Tropical and geographical medicine. 1968;20(3):251–6.

11. Navarrete-Espinosa J, Gomez-Dantes H. Arbovirus causales de fiebre hemorrágica en pacientes del Instituto Mexicano del Seguro Social. Revista medica del Instituto Mexicano del Seguro Social. 2006;44(4):347–53.

12. Groot H. Estudios sobre virus transmitidos por artropodos en Colombia. Rev Acad Colomb Cienc Exactas Fis Nat. 1964;12(46):191–217.

13. Talarmin A, Chandler LJ, Kazanji M, de Thoisy B, Debon P, Lelarge J, et al. Mayaro virus fever in French Guiana: isolation, identification, and seroprevalence. The American journal of tropical medicine and hygiene. 1998;59(3):452–6.

14. Blohm G, Elbadry MA, Mavian C, Stephenson C, Loeb J, White S, et al. Mayaro as a Caribbean traveler: Evidence for multiple introductions and transmission of the virus into Haiti. International journal of infectious diseases : IJID : official publication of the International Society for Infectious Diseases. 2019;87:151–3.

15. Hoch AL, Peterson NE, LeDuc JW, Pinheiro FP. An outbreak of Mayaro virus disease in Belterra, Brazil. III. Entomological and ecological studies. The American journal of tropical medicine and hygiene. 1981;30(3):689–98.

16. Peterson AT. Ecologic niche modeling and spatial patterns of disease transmission. Emerging infectious diseases. 2006;12(12):1822–6.

17. Ochieng AO, Nanyingi M, Kipruto E, Ondiba IM, Amimo FA, Oludhe C, et al. Ecological niche modelling of Rift Valley fever virus vectors in Baringo, Kenya. Infection ecology & epidemiology. 2016;6:32322.

18. Gurgel-Gonçalves R, Galvão C, Costa J, Peterson AT. Geographic distribution of chagas disease vectors in Brazil based on ecological niche modeling. J Trop Med. 2012;2012:705326.

19. Miller RH, Masuoka P, Klein TA, Kim HC, Somer T, Grieco J. Ecological niche modeling to estimate the distribution of Japanese encephalitis virus in Asia. PLoS neglected tropical diseases. 2012;6(6):e1678.

20. Lorenz C, Freitas Ribeiro A, Chiaravalloti-Neto F. Mayaro virus distribution in South America. Acta tropica. 2019;198:105093.

21. de Almeida MA, Dos Santos E, Cardoso JdC, da Silva LG, Rabelo RM, Bicca-Marques JC. Predicting yellow fever through species distribution modeling of virus, vector, and monkeys. EcoHealth. 2019;16(1):95–108.

22. GBIF.org (24 December 2020) GBIF Occurrence Download https://doi.org/10.15468/dl.jue5tw.

23. Benson DA, Cavanaugh M, Clark K, Karsch-Mizrachi I, Lipman DJ, Ostell J, et al. GenBank. Nucleic Acids Research. 2016;45(D1):D37–D42.

24. Foley DH, Wilkerson RC, Rueda LM. Importance of the “what,” “when,” and “where” of mosquito collection events. Journal of medical entomology. 2009;46(4):717–22.

25. Wieczorek J, Guo Q, Hijmans R. The point-radius method for georeferencing locality descriptions and calculating associated uncertainty. International journal of geographical information science. 2004;18(8):745–67.

26. Foley DH, Pecor DB. Best Practices Guide to Entomological Surveillance: Data Management and Reporting. : WRBU Information Products; 2016 [Available from: http://vectormap.si.edu/.

27. Phillips SJ, Anderson RP, Schapire RE. Maximum entropy modeling of species geographic distributions. Ecological Modelling. 2006;190:231–59.

28. Merow C, Smith, M.J., Silander, J.A. A practical guide to MaxEnt for modeling species’ distributions: what it does, and why inputs and settings matter. Ecography. 2013;36:1058–69.

29. Hernandez PA, Graham CG, Master LL, Albert DL. The effect of sample size and species characteristics on performance of different species distribution modeling methods. Ecography. 2006;29(5):773–85.

30. Elith J. Novel methods improve prediction ofspecies’ distributions from occurrence data. Ecography. 2006;29:129–51.

31. Fick SE, Hijmans RJ. WorldClim 2: new 1km spatial resolution climate surfaces for global land areas. International Journal of Climatology. 2017;37(12):4302–15.

32. Danielson JJ, Gesch DB. Global Multi-resolution Terrain Elevation Data 2010 (GMTED2010). 2011.

33. ESRI. World Land Cover at 30m resolution from MDAUS BaseVue 2013. 2015.

34. ESRI. Soils of the world from the from the United Nations Food and Agriculture Organization. 2014.

35. ESRI. ArcGIS Desktop: Release 10.8 Redlands, CA: Environmental Systems Research Institute 2019.

36. Runfola D, Anderson A, Baier H, Crittenden M, Dowker E, Fuhrig S, et al. geoBoundaries: A global database of political administrative boundaries. PloS one. 2020;15(4):e0231866.

37. Zar JH. Biostatistical Analysis. Upper Saddle River, New Jersey: Prentice Hall:. 663 pp.; 1999. 663 p.

38. Phillips SJ, Dudík M, Elith J, Graham CH, Lehmann A, Leathwick J, et al. Sample selection bias and presence-only distribution models: implications for background and pseudo-absence data. Ecol Appl. 2009;19(1):181–97.

39. Kramer-Schadt S, Niedballa J, Pilgrim JD, Schröder B, Lindenborn J, Reinfelder V, et al. The importance of correcting for sampling bias in MaxEnt species distribution models. Diversity and Distributions. 2013;19(11):1366–79.

40. Phillips SJ, Anderson RP, Dudik M, Schapire RE, Blair ME. Opening the black box: an open-source release of Maxent. Ecography. 2017;40:887–93.

41. Richman R, Diallo D, Diallo M, Sall AA, Faye O, Diagne CT, et al. Ecological niche modeling of Aedes mosquito vectors of chikungunya virus in southeastern Senegal. Parasites & vectors. 2018;11(1):255.

42. Warren DL, Seifert, S.N. Ecological niche modeling in Maxent: the importance of model complexity and the performance of model selection criteria. Ecological Applications. 2011;21(2):335–42.

43. Liria J, Navarro J-C. Modelo de nicho ecológico en Haemagogus Williston (Diptera: Culicidae), vectores del virus de la fiebre amarilla. Revista Biomédica. 2010;21(3):149–61.

44. Pan American Health Organization / World Health Organization. Epidemiological Alert: Mayaro Fever. Washington, D.C.: PAHO/WHO; 2019 May 1, 2019.

45. Hozé N, Salje H, Rousset D, Fritzell C, Vanhomwegen J, Bailly S, et al. Reconstructing Mayaro virus circulation in French Guiana shows frequent spillovers. Nat Commun. 2020;11(1):2842.

46. World Health Organization. Mayaro virus disease - French Guiana, France 2020 [Available from: https://www.who.int/csr/don/25-october-2020-mayaro-fever-french-guiana-france/en/.

47. Causey OR, Maroja OM. Mayaro virus: a new human disease agent. III. Investigation of an epidemic of acute febrile illness on the river Guama in Para, Brazil, and isolation of Mayaro virus as causative agent. The American journal of tropical medicine and hygiene. 1957;6(6):1017–23.

48. LeDuc JW, Pinheiro FP, Travassos da Rosa AP. An outbreak of Mayaro virus disease in Belterra, Brazil. II. Epidemiology. The American journal of tropical medicine and hygiene. 1981;30(3):682–8.

49. Auguste AJ, Liria J, Forrester NL, Giambalvo D, Moncada M, Long KC, et al. Evolutionary and Ecological Characterization of Mayaro Virus Strains Isolated during an Outbreak, Venezuela, 2010. Emerging infectious diseases. 2015;21(10):1742–50.

50. Silva NIO, Sacchetto L, de Rezende IM, Trindade GS, LaBeaud AD, de Thoisy B, et al. Recent sylvatic yellow fever virus transmission in Brazil: the news from an old disease. Virology journal. 2020;17(1):9.

51. Ali R, Mohammed A, Jayaraman J, Nandram N, Feng RS, Lezcano RD, et al. Changing patterns in the distribution of the Mayaro virus vector Haemagogus species in Trinidad, West Indies. Acta tropica. 2019;199:105108.

52. Alencar J, Mello CF, Morone F, Albuquerque HG, Serra-Freire NM, Gleiser RM, et al. Distribution of Haemagogus and Sabethes Species in Relation to Forest Cover and Climatic Factors in the Chapada Dos Guimarães National Park, State of Mato Grosso, Brazil. Journal of the American Mosquito Control Association. 2018;34(2):85–92.

53. Izurieta RO, Macaluso M, Watts DM, Tesh RB, Guerra B, Cruz LM, et al. Hunting in the Rainforest and Mayaro Virus Infection: An emerging Alphavirus in Ecuador. Journal of global infectious diseases. 2011;3(4):317–23.

54. Abad-Franch F, Grimmer GH, de Paula VS, Figueiredo LT, Braga WS, Luz SL. Mayaro virus infection in amazonia: a multimodel inference approach to risk factor assessment. PLoS neglected tropical diseases. 2012;6(10):e1846.

55. Tesh RB, Watts DM, Russell KL, Damodaran C, Calampa C, Cabezas C, et al. Mayaro virus disease: an emerging mosquito-borne zoonosis in tropical South America. Clinical infectious diseases : an official publication of the Infectious Diseases Society of America. 1999;28(1):67–73.

56. Black FL, Hierholzer WJ, Pinheiro F, Evans AS, Woodall JP, Opton EM, et al. Evidence for persistance of infectious agents in isolated human populations. Am J Epidemiol. 1974;100(3):230–50.

57. Beck-Johnson LM, Nelson WA, Paaijmans KP, Read AF, Thomas MB, Bjørnstad ON. The importance of temperature fluctuations in understanding mosquito population dynamics and malaria risk. R Soc Open Sci. 2017;4(3):160969.

58. Carrington LB, Seifert SN, Willits NH, Lambrechts L, Scott TW. Large diurnal temperature fluctuations negatively influence Aedes aegypti (Diptera: Culicidae) life-history traits. Journal of medical entomology. 2013;50(1):43–51.

59. Lambrechts L, Paaijmans KP, Fansiri T, Carrington LB, Kramer LD, Thomas MB, et al. Impact of daily temperature fluctuations on dengue virus transmission by Aedes aegypti. Proceedings of the National Academy of Sciences of the United States of America. 2011;108(18):7460–5.

60. Chadee DD, Tikasingh ES, Ganesh R. Seasonality, biting cycle and parity of the yellow fever vector mosquito Haemagogus janthinomys in Trinidad. Medical and veterinary entomology. 1992;6(2):143–8.

