## Supplemental Table 2 for "An ecological niche model to predict the geographic distribution of *Haemagogus janthinomys*, Dyar, 1921 the yellow fever and Mayaro virus vector, in South America"

**S2 Table.** **Minimum, maximum, average values, percent contribution, and permutation importance of variables in the full *Hg. janthinomys* model**

| **Variable** | **Description** | | **Min.** | **Max.** | **Avg.** | **Contribution (%)** | **Permutation (%)** |
| --- | --- | --- | --- | --- | --- | --- | --- |
| **Land Use/Land Cover** | Categorical variable with 16 classes of land use / land cover | | N/A^a^ | N/A | N/A | 32.4 | 6.2 |
| **BIO2** | Mean Diurnal Range (Mean of monthly (max temp - min temp)), °C | | 6.3 | 14.0 | 9.4 | 25.3 | 10.5 |
| **BIO5** | Max Temperature of Warmest Month, °C | | 25.4 | 34.2 | 30.8 | 7.6 | 8 |
| **BIO4** | Temperature Seasonality (standard deviation ×100) | | 37.3 | 587.0 | 106.5 | 5.1 | 13.4 |
| **Aspect** | The direction of each cell’s slope, measured in degrees (from 0 to 360) | | 16.6 | 358.9 | 172.4 | 4.9 | 2.2 |
| **Soils** | Categorical variable that defines 106 soil units and 4 non-soil units | | N/A^a^ | N/A | N/A | 4.3 | 1.5 |
| **Slope** | The angle of the downward sloping terrain from 0 (flat) to 90 degrees (vertical) | | 0 | 33 | 4.9 | 4.3 | 2.4 |
| **Elevation** | Digital elevation model, m | | 0 | 1328 | 308.8 | 2.8 | 6.6 |
| **BIO18** | Precipitation of Warmest Quarter, mm | | 67 | 869 | 421.1 | 2.2 | 4.1 |
| **BIO13** | Precipitation of Wettest Month, mm | | 81 | 582 | 304.8 | 2.0 | 2.7 |
| **BIO14** | Precipitation of Driest Month, mm | | 1 | 231 | 58.3 | 1.6 | 4.8 |
| **BIO19** | Precipitation of Coldest Quarter, mm | | 6 | 1312 | 496.7 | 1.6 | 1.8 |
| **Flow Direction** | Flow direction from each cell to steepest downslope neighbor. Eight possible values represent direction of flow | | N/A^a^ | N/A | N/A | 1.5 | 0.8 |
| **BIO17** | Precipitation of Driest Quarter, mm | | 6 | 724 | 203.2 | 0.9 | 9.7 |
| **BIO9** | Mean Temperature of Driest Quarter, °C | | 11.4 | 28.1 | 23.7 | 0.7 | 6.4 |
| **BIO15** | Precipitation Seasonality (Coefficient of Variation), mm | | 10.2 | 114.8 | 55.5 | 0.6 | 2.8 |
| **Flow Accumulation** | Accumulated weight of all cells in the Flow Direction raster flowing into each downslope cell | | 0 | 20 | 1.9 | 0.5 | 1.0 |
| **BIO3** | Isothermality (BIO2/BIO7) (×100), °C | | 46.3 | 90.0 | 73.8 | 0.5 | 0.9 |
| **BIO16** | Precipitation of Wettest Quarter, mm | | 213 | 1414 | 824.3 | 0.5 | 4.2 |
| **BIO7** | Temperature Annual Range (BIO5-BIO6), °C | | 8.8 | 30.3 | 13.1 | 0.2 | 3.3 |
| **BIO11** | Mean Temperature of Coldest Quarter, °C | | 11.4 | 26.7 | 23.0 | 0.1 | 0.2 |
| **BIO12** | Annual Precipitation, mm | | 363 | 3695 | 2002.9 | 0.1 | 4.0 |
| **BIO8** | Mean Temperature of Wettest Quarter, °C | | 19.6 | 27.5 | 24.8 | 0.1 | 0.5 |
| **BIO6** | Min Temperature of Coldest Month, °C | | 2.9 | 23.1 | 17.7 | 0.1 | 1.2 |
| **BIO10** | Mean Temperature of Warmest Quarter, °C | | 19.6 | 28.1 | 25.6 | 0 | 0.6 |
| **BIO1** | Annual Mean Temperature, °C | | 16.8 | 27.1 | 24.4 | 0 | 0.1 |
| ^a^ Minimum, maximum, and average values were not calculated for categorical variables. | | | | | | | |
